# A catalogue of small proteins from the global microbiome

**DOI:** 10.1101/2023.12.27.573469

**Authors:** Yiqian Duan, Celio Dias Santos-Junior, Thomas Sebastian Schmidt, Anthony Fullam, Breno L. S. de Almeida, Chengkai Zhu, Kuhn Michael, Xing-Ming Zhao, Peer Bork, Luis Pedro Coelho

**Affiliations:** Institute of Science and Technology for Brain-Inspired Intelligence, Fudan University, Shanghai, China; Laboratory of Microbial Processes & Biodiversity - LMPB; Department of Hydrobiology, Universidade Federal de São Carlos – UFSCar, São Carlos, São Paulo, Brazil; Structural and Computational Biology Unit, European Molecular Biology Laboratory, Heidelberg, Germany; Department of Neurology, Zhongshan Hospital, Fudan University, Shanghai, China; State Key Laboratory of Medical Neurobiology, Institutes of Brain Science, Fudan University, Shanghai, China; MOE Key Laboratory of Computational Neuroscience and Brain-Inspired Intelligence; MOE Frontiers Center for Brain Science, Fudan University, Shanghai, China; International Human Phenome Institute, Shanghai, China; Max Delbrück Centre for Molecular Medicine, Berlin, Germany; Department of Bioinformatics, Biocenter, University of Würzburg, Würzburg, Germany; Centre for Microbiome Research, School of Biomedical Sciences, Queensland University of Technology, Translational Research Institute, Woolloongabba, Queensland, Australia

## Abstract

Small open reading frames (smORFs) shorter than 100 codons are widespread and perform essential roles in microorganisms, where they encode proteins active in several cell functions, including signal pathways, stress response, and antibacterial activities. However, the ecology, distribution and role of small proteins in the global microbiome remain unknown. Here, we constructed a global microbial smORFs catalogue (GMSC) derived from 63,410 publicly available metagenomes across 75 distinct habitats and 87,920 high-quality isolate genomes. GMSC contains 965 million non-redundant smORFs with comprehensive annotations. We found that archaea harbor more small proteins proportionally than bacteria. We moreover provide a tool called GMSC-mapper to identify and annotate small proteins from microbial (meta)genomes. Overall, this publicly-available resource demonstrates the immense and underexplored diversity of small proteins.

## Introduction

Small open reading frames (smORFs) are found in all three domains of life, estimated as 5-10% of annotated genes^1–3^. Small proteins encoded by smORFs have been reported to perform key functions in microbial cells^4–8^ and have been found involved in transcription to regulate gene expression^9^, to stabilize large protein complexes^10^, in signaling transduction pathways^11^, regulation of transporters^12^, sporulation^13,14^, photosynthesis^15^, and response to environmental cues^16^. In addition, small proteins can also perform antibacterial activities^17^ or compose toxin/antitoxin (TA) systems^18,19^.

However, small proteins have been neglected in (meta)genomics-based global studies of the microbiome^20,21^ due to the difficulty in reliably identifying smORFs using genomic information alone^22,23^. Advances in Ribo-Seq^24^ and proteogenomics methods^25,26^ combined with comparative genomics methods^27,28^ have enabled the discovery of an increasing number of small proteins in various microorganisms^29–32^. For example, a recent systematic study revealed 4,539 novel conserved small protein families of the human microbiome^33^, 30% of which are predicted to encode transmembrane or secreted proteins. However, most of the studies focusing on smORFs approach isolated microorganisms and specific environments. The functional and ecological understanding of microbial smORFs at a global scale across different habitats is still very limited.

Here, we used the principle that repeated independent observations of the same small protein (or minor variations thereof) minimize the likelihood of false positive smORF predictions and constructed a global microbial smORFs catalogue (GMSC) derived from 63,410 assembled metagenomes from the SPIRE database^21^ and 87,920 isolate genomes from the ProGenomes2 database^34^. In the catalogue, we provide comprehensive annotation containing taxonomy classification, habitat assignment, quality assessment, conserved domain annotation, and predicted cellular localization.

In addition, the GMSC can be used as a reference to annotate (meta)genomes as the presence of homologues reduces the probability that false positives are reported. To facilitate this, we developed a tool, named GMSC-mapper, which additionally provides users with information about the distribution of any matching smORFs across taxonomy, habitats, and geography. Thus, our catalogue and associated tools can be used to study the presence, prevalence, distribution, and potential ecological roles of smORFs on a global scale, and provide new insights into how these molecules work within microorganisms.

## Results

### The global microbial smORFs catalogue comprises 965 million small proteins

The global microbial smORFs catalogue (GMSC) was derived from 63,410 publicly available assembled metagenomes spanning multiple habitats worldwide from the SPIRE database^21^ and 87,920 high-quality isolate microbial genomes from the ProGenomes2 database^34^ (Fig. 1a, Supplementary Table 1). From the assembled contigs, we used the modified version of Prodigal^35^ in Macrel^36^ to predict open reading frames (ORFs) with a minimum length of 30 nucleotides (see Methods). The ORFs encoding small proteins (here defined as those up to 100 amino acids) were considered small ORFs (smORFs).

**Fig. 1.**
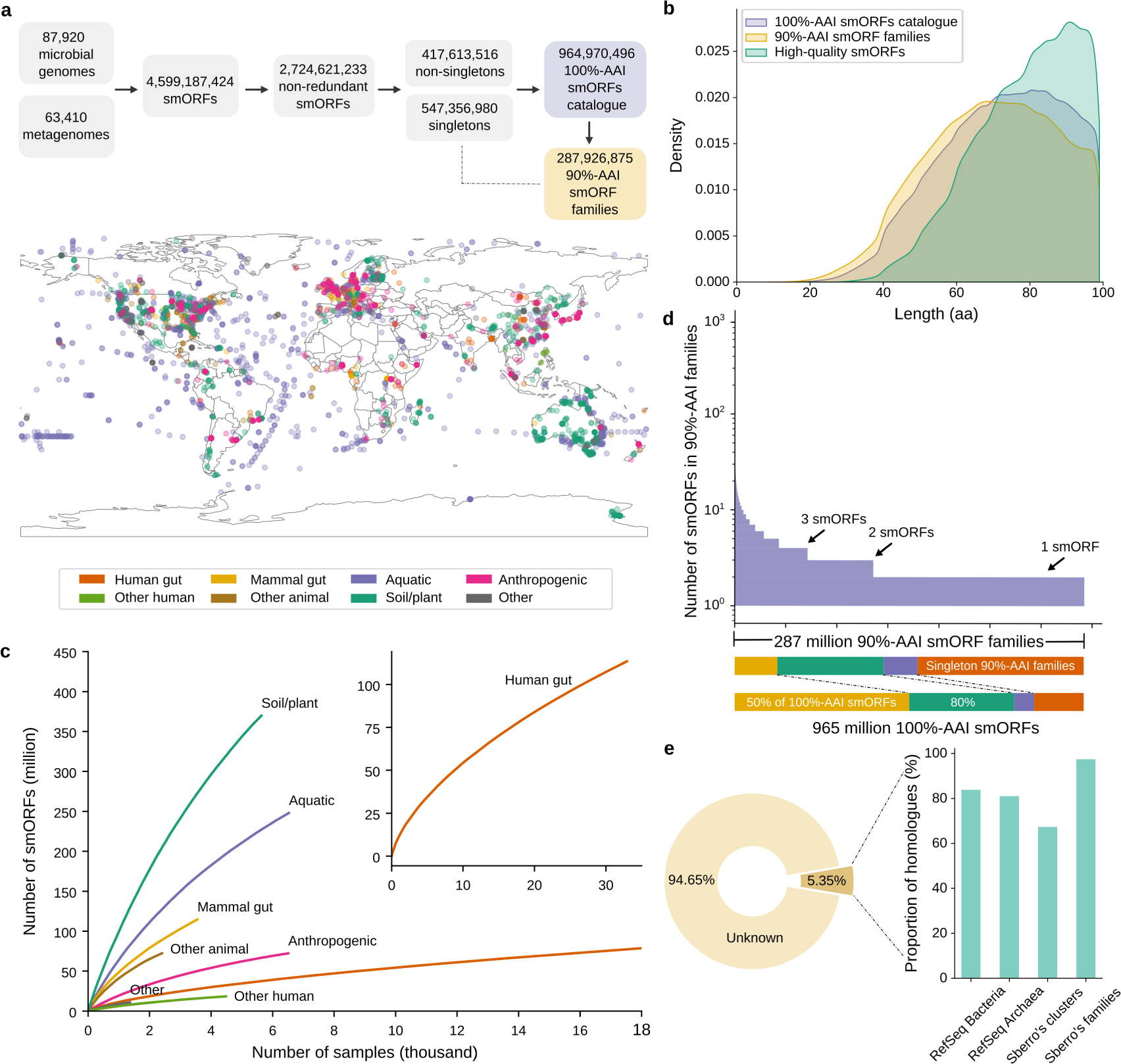
Global Microbial smORFs Catalogue (GMSC) **(a)** ORFs (Open Reading Frames) were predicted from contigs from 63,410 publicly available metagenomes and 87,920 microbial genomes. The ORFs with at most 300 bps were considered smORFs. We hierarchically clustered non-redundant smORFs into clusters that are 90% amino acid identical (Methods). **(b)** Small proteins encoded by smORFs range in length from 9 to 99 amino acids. Sequences that pass all *in silico* quality tests and contain at least one piece of experimental evidence are considered *high-quality predictions* (Methods). **(c)** Shown are gene accumulation curves per habitat, showing how sampling affects the discovery of smORFs (see also Supplementary Fig. 2a). **(d)** The largest 90%-AAI smORF family contains 4,577 sequences. The size of 90%-AAI smORF families exhibits a long tail distribution and 47.5% of families consist of only one sequence, accounting for fewer than 15% of the total GMSC smORFs. A small fraction of large families account for the majority of GMSC smORFs (12.2% of families contain 50% of smORFs). **(e)** Only 5.35% of smORFs in the GMSC have a homologous sequence in another sequence catalogue (Methods). On the other hand, more than 80% of bacterial and archaeal small proteins from the RefSeq database have a homologue in our catalogue. Although only 67.3% of the 444,054 small protein clusters from the Sberro human microbiome dataset are homologous to a protein in our catalogue, most of their clusters without homologous sequences only contain one sequence. Among the 4,539 conserved small protein families from the Sberro human microbiome dataset, 97.4% of them are homologous to our catalogue.

In total, after collapsing smORFs coding for identical amino acid sequences, we obtained 2,724,621,233 smORFs. A large majority (84.7%) was of singleton sequences. To reduce the incidence of false positives^36^, we focused first on the 417 million non-singleton sequences. We hierarchically clustered these non-singleton smORFs at 90% amino acid identity and 90% coverage, which resulted in 287,926,875 clusters, which we will henceforth refer to as *families*. Then, we constructed the *smORFs catalogue*, which contains both non-singletons as well as any singleton that matches a family representative at 90% amino acid identity and 90% coverage (*rescued singletons*, see Methods). The final *smORFs catalogue* contains 964,970,496 smORFs.

The samples in our dataset had been previously manually curated into 75 habitats^21^, which we further grouped into 8 broad categories: mammal gut, anthropogenic, other-human, other-animal, aquatic, human gut, soil/plant, and other (Methods, Supplementary Table 2). Despite the large number of samples we have collected, rarefaction analysis indicates that smORF diversity is far from covered (Fig. 1c; Supplementary Fig. 2a).

Approximately half of GMSC families consist of only one sequence, but the size distribution of families is long-tailed, so that the largest 12.2% of families already cover half of the 100AA smORFs (Fig. 1d).

### 43 million smORFs are high-quality and most smORFs represent novelty

Predicting smORFs can result in a high rate of false positives. Thus, in addition to discarding non-homologue singleton predictions, we performed several *in silico* quality tests including estimating coding potential of families using RNAcode^37^ and additionally matching genomic predictions to publicly-available metatranscriptomic and metaproteomics data (see Methods). In total, 43,617,914 (4.5%) of the smORFs pass all *in silico* quality tests and have at least one match in transcriptional or translational data. We henceforth refer to these as *high-quality predictions* (Supplementary Fig. 3a-c).

To assess the novelty and comprehensiveness of our catalogue, we matched small proteins encoded by GMSC smORFs to the RefSeq database^38^ and previously published human microbiome small protein family datasets^33^. Only 5.3% of smORFs in our catalogue are homologous to these previously reported small proteins (Fig. 1e). On the other hand, our catalogue contains more than 80% of these reference datasets. For smORFs of *high-quality predictions*, a higher proportion (8.7%) show homology with these reference datasets, but they only cover *circa* 20% of the reference datasets (Supplementary Fig. 4a). Hence the *high-quality predictions* produce a large number of novel small proteins with high confidence that are not present in other reference datasets, but as the available transcriptome and metaproteome datasets are limited, discarding non-high-quality predictions would result in a large loss of coverage.

To explore the functions undertaken by the small proteins encoded by the smORFs in our catalogue, we searched the small protein families against the Conserved Domain Database (CDD)^39^ using RPS-BLAST^40,41^. Only 6.1% of small protein families containing 86,694,259 smORFs (8.98%) were assigned CDD domains, compared to 35.2% of canonical-length proteins (greater than 100 amino acids)^20^. As expected, smORFs in *high-quality predictions* are twice as likely to be assigned a CDD domain (18.8%, P-value < 10*^−^* ^308^, hypergeometric test).

### Even conserved small proteins lack functional annotations

Using MMSeqs2 taxonomy^42^ we predicted the taxonomic origin of contigs and transferred that prediction to the smORFs (Methods). This process returned a prediction for 81.6% of the 100AA smORFs, with more than half (56.9%) being assigned to a genus or species (Fig. 2a).

**Fig. 2.**
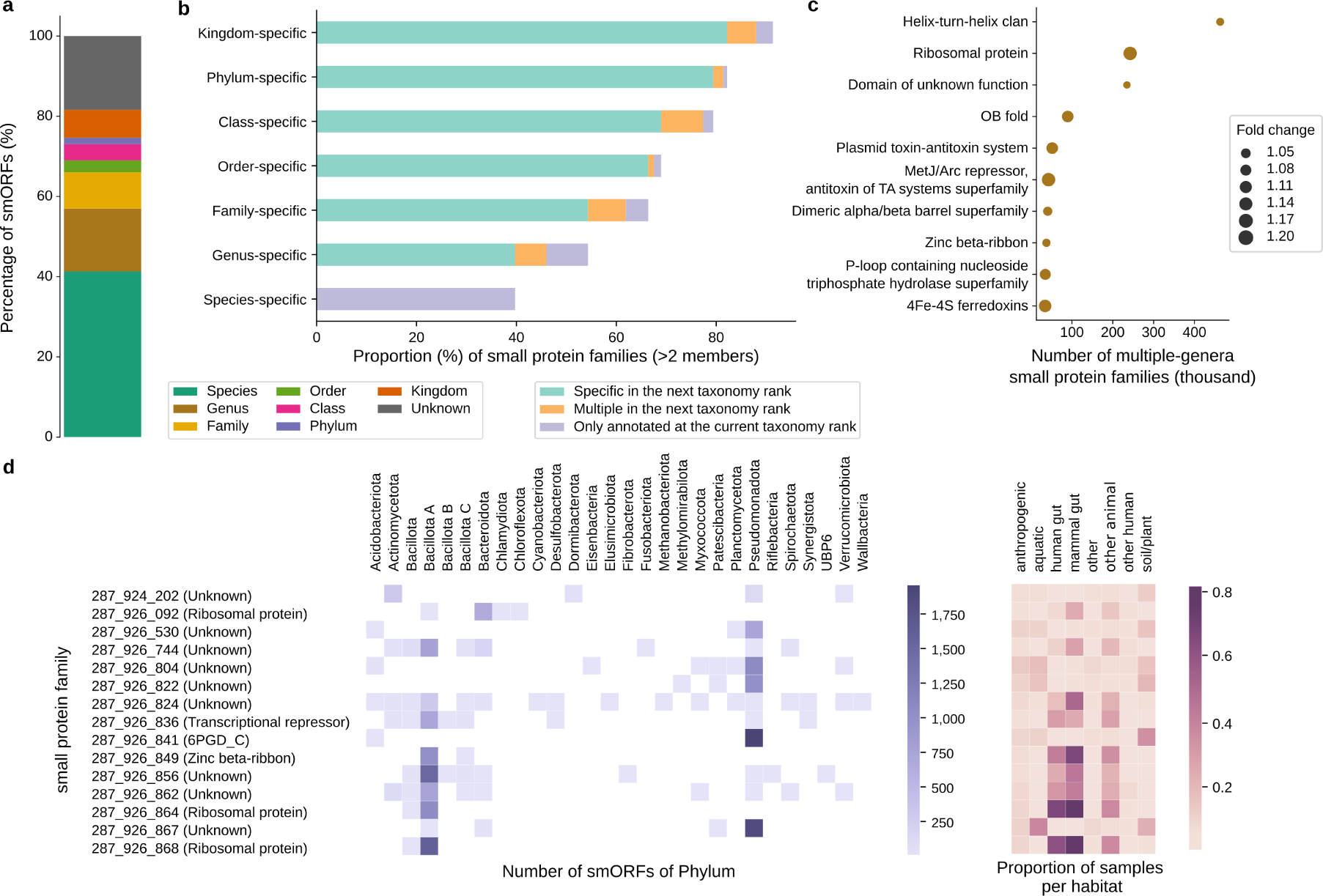
Taxonomic and functional annotation of small proteins. **(a)** Predicting taxonomy for the contigs and genomes from which smORFs originate (Methods) resulted in a taxonomic assignment for 81.6% of smORFs (56.9% of smORFs at genus or species level). **(b)** When only families with >2 members were considered (96,721,815 families), there are three cases at each taxonomic rank. For example, considering the rank of *class*, a small protein family is annotated to a particular taxonomic class if all its members are annotated as belonging to that class (unannotated smORFs being ignored). We further distinguish three cases, namely whether its members are (i, marked *specific in the next taxonomic rank*) all be annotated to the same order (as order is the next taxonomic rank), (ii, marked *multiple in the next taxonomic rank*) annotated to different orders within that class, or (iii, marked *only annotated at the current taxonomic rank*) not annotated to any order. Other ranks are treated analogously (until we reach the level of species). **(c)** The enrichment of Pfam domains in small protein families present in multiple genera compared to the entire families with over two members (P-value < 0.05, hypergeometric test, corrected by Bonferroni). Pfam domains were grouped by Pfam domain clans. Fold change is the ratio of the Pfam proportion of small protein families which present in multiple genera to the Pfam proportion of the entire families with over two members. **(d)** The Pfam annotation of small protein families that exist in multiple phyla, spanning >100 species and distributed across all the 8 broad habitat categories (mammal gut, anthropogenic, other-human, other-animal, aquatic, human gut, soil/plant, and other).

We next investigated the taxonomic breadth and conservation of smORFs^28,43^. Of the 96,721,815 small protein families with at least three members, more than half of them (52,550,829) are genus-specific (Fig. 2b). Among these genus-specific families, most are species-specific, accounting for 39.7% of the families included in the analysis.

We reasoned that the multi-genus families may be especially likely to be present in multiple habitats and involved in critical cellular functions^33^. As expected, multi-genus families are more common in multiple habitats than the entire set of families with at least three members even when differences in family size distributions are taken into account, but the difference is not large (61.8% vs. 57.5%; P-value < 10*^−^* ^308^, due to the large number of datapoints, hypergeometric test). Furthermore, we traced the conserved Pfam domains of small protein families^44^ (Supplementary Table 4). Multi-genus families are annotated with Pfam domains at a higher rate than the background of all families with at least three members (9.91% vs 8.15%; P-value < 10*^−^* ^308^, hypergeometric test). Nonetheless, it is noteworthy that the vast majority have no detected Pfam domain and that a further 9.5% of those annotated, were annotated with Pfam domains of unknown functions (Fig. 2c).

We then focused on conserved families present in multiple phyla. We found a total of 2,437 multi-phylum families present across all 8 broad habitat categories (Supplementary Table 5). Of these, only 752 families were annotated with Pfam conserved domains, of which 268 (35.6%) were associated with ribosomal proteins and 99 (13.2%) belonged to the Helix-turn-helix clan (Fig. 2d).

### Archaea harbor more smORFs proportionally than bacteria

To investigate the presence of smORFs in different microorganisms without sampling bias, we calculated the number of redundant smORFs per megabase pairs (Mbp) of assembled contigs, also named as the density of smORFs^32,45^.

Most of the genera with the highest density come from *Pseudomonadota*, *Bacillota A*, *Actinomycetota*, *Bacillota*, and *Bacteroidota* (Fig. 3a). However, when considering the density of phyla as a whole, interestingly, we found the density of archaeal phyla is higher than bacterial ones (*P_Mann_* = 2.25 *·* 10*^−^*^3^; Fig. 3b). Of the ten phyla with the highest smORF density, half are archaeal, despite the fact that only 18 archaeal phyla contained enough data to be analysed compared to 131 bacterial ones (Fig. 3c, Supplementary Table 6). The phyla that produce the most smORFs per Mbp are *Desulfobacterota D* (362.87 smORFs per Mbp), *Undinarchaeota* (331.35 smORFs per Mbp), *Nanoarchaeota* (281.34 smORFs per Mbp), *Methylomirabilota* (241.37 smORFs per Mbp), and *Huberarchaeota* (241.05 smORFs per Mbp).

**Fig. 3.**
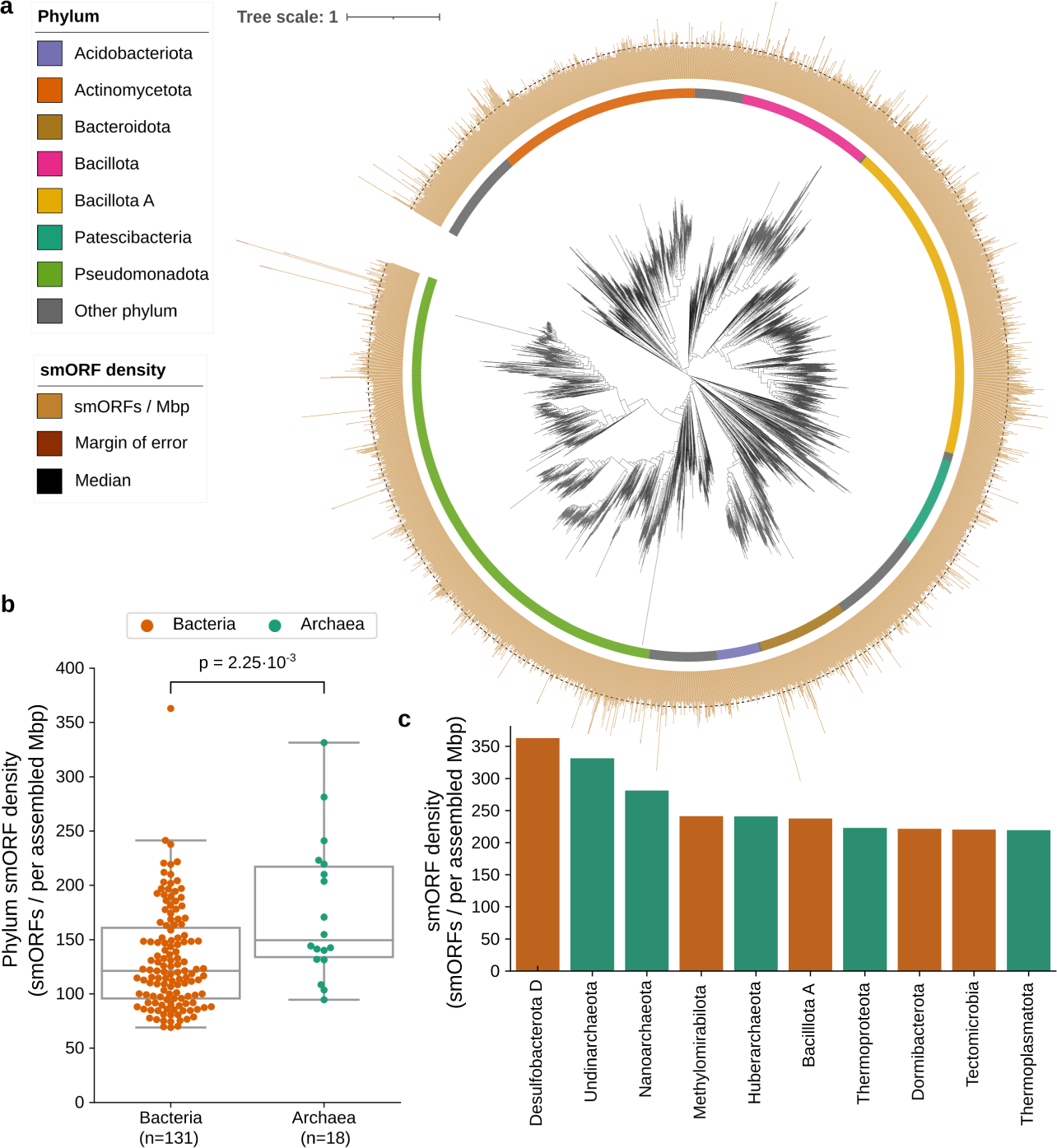
Archaea harbor more smORFs than bacteria. **(a)** Shown are the smORFs density distribution for the top 3,000 bacterial genera with the highest density (brown bars, confidence interval of 95% shown as dark brown bars). Most of the densest genera are from *Pseudomonadota*, *Bacillota A*, and *Actinomycetota*. For reference, the black dashed line represents the median smORFs density for the presented genera. **(b)** Calculating the smORFs density of each phylum, the density of archaea is significantly higher than that of bacteria. **(c)** The top 10 phyla with the highest smORF density are shown.

### Archaea have more transmembrane or secreted small proteins than bacteria

Given the higher densities of smORFs in Archaea, we investigated the functions and properties in archaeal and bacterial small proteins encoded by smORFs^46^. We compared the archaeal and bacterial small protein families with COG^47^ annotation. Only 1.72% of the families are annotated with COGs, of which 4,747,223 families are from bacteria and 202,825 families are from archaea. The COG classes that belong to Information storage and processing account for the largest proportion of small proteins in both bacteria and archaea (Fig. 4a), which is consistent with that found by Wang et al^43^. However, *circa* 17% of small proteins in bacteria and archaea are still annotated as COG classes which are poorly characterized.

**Fig. 4.**
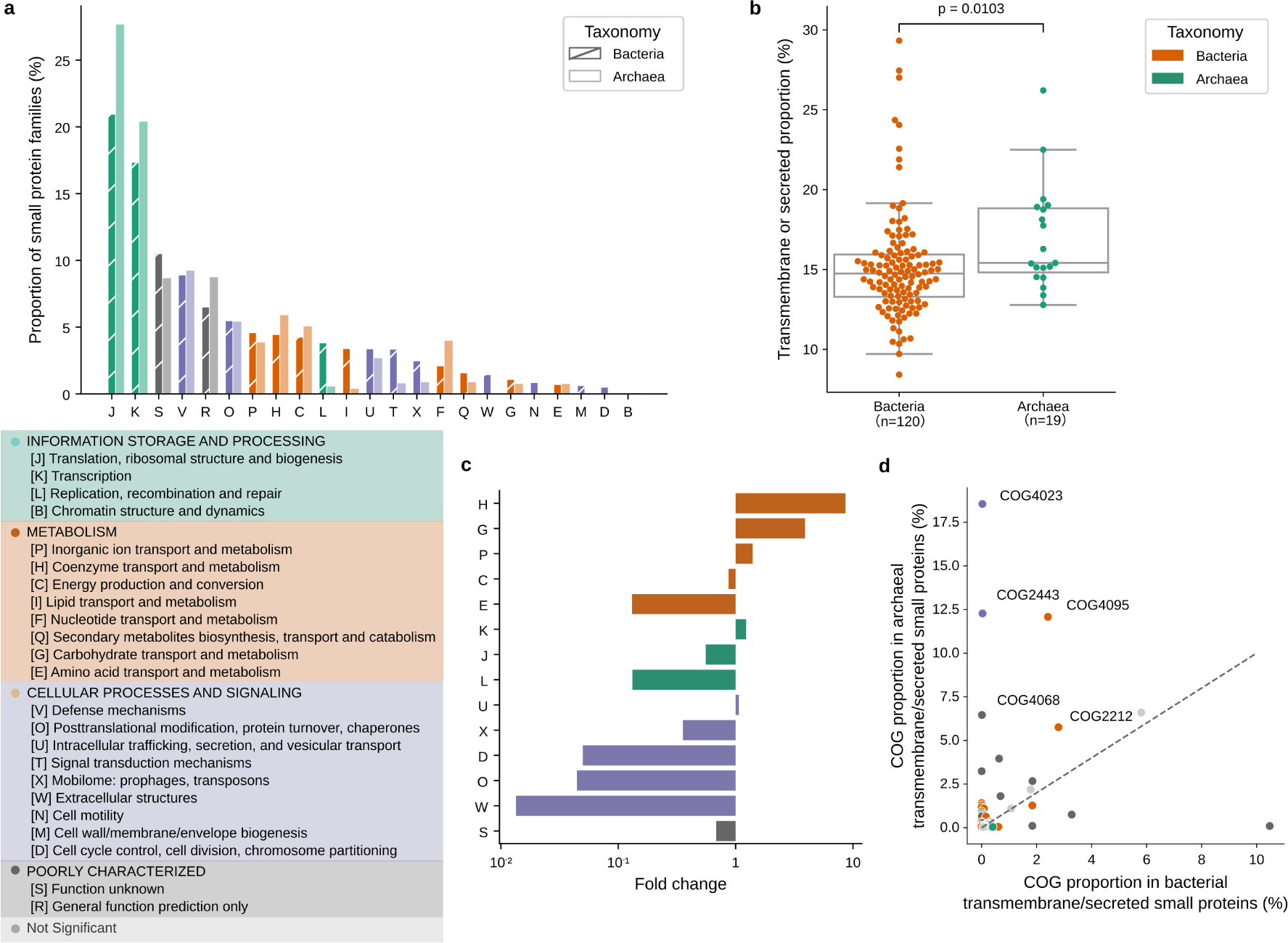
Archaea have more transmembrane or secreted small proteins than bacteria. **(a)** The COG distribution of archaeal and bacterial small proteins is shown. **(b)** Archaea contain a higher fraction of transmembrane or secreted small proteins than bacteria (calculated per phylum). P-values shown are from the Mann-Whitney Test (two-sided) with multiple tests adjusted by Bonferroni correction. **(c)** Shown is the difference in the proportion of COG class in archaeal transmembrane or secreted small proteins versus bacterial transmembrane or secreted small proteins. The fold change is the ratio of proportions. The p-values were calculated using Fisher’s exact test and adjusted by Bonferroni correction. **(d)** Dots represent 43 COGs, which are enriched in archaeal transmembrane or secreted small proteins compared to the archaeal small proteins that are not transmembrane or secreted as well as bacterial transmembrane or secreted small proteins. The proportion comparison of these 43 COGs between archaeal transmembrane or secreted small proteins and bacterial transmembrane or secreted small proteins is shown.

Small proteins with transmembrane or secreted characteristics may be involved in cell communication^7^. We explored the transmembrane and secreted small proteins in archaea and bacteria (Methods). 15.3% of the families are predicted to be potentially transmembrane (using TMHMM-2.0^48^) or secreted (using SignalP-5.0^49^). Archaeal small proteins are more likely than bacterial ones to be transmembrane or secreted (*P_M a nn_* ≤ 0.0103, Fig. 4b)^50^.

Furthermore, compared with bacterial transmembrane or secreted small proteins, we found that archaeal transmembrane or secreted small proteins are enriched in COG classes related to the transport and metabolism of coenzymes, carbohydrates, and inorganic ions, besides the intracellular trafficking, secretion, and vesicular transport. In contrast, they are depleted in COG classes related to cellular processes and signaling (P-value < 0.05, Fisher’s exact test, multiple tests corrected by Bonferroni, Fig. 4c).

Some COGs were primarily (or even exclusively) present in archaea (as defined by a P-value < 0.05, Fisher’s exact test, multiple tests corrected by Bonferroni, Fig. 4d). For example, the COG with the highest proportion in archaea, COG4023 is a preprotein translocase subunit Sec61beta, which is a component of the Sec61/SecYEG protein secretion system. It is found in eukaryotes and archaea and is possibly homologous to the bacterial SecG^51^.

### Identification of smORFs by GMSC-mapper

As mentioned above, smORF predictions are prone to false positives and one strategy for increasing confidence is to find sequences present in multiple genomes (or metagenomes). In this context, our catalogue can be a resource whereby users with a single sample (or a small number of samples) use it as a reference to obtain high-quality predicted smORFs. For this usage, we provide a tool called GMSC-mapper (Fig. 5a).

**Fig. 5.**
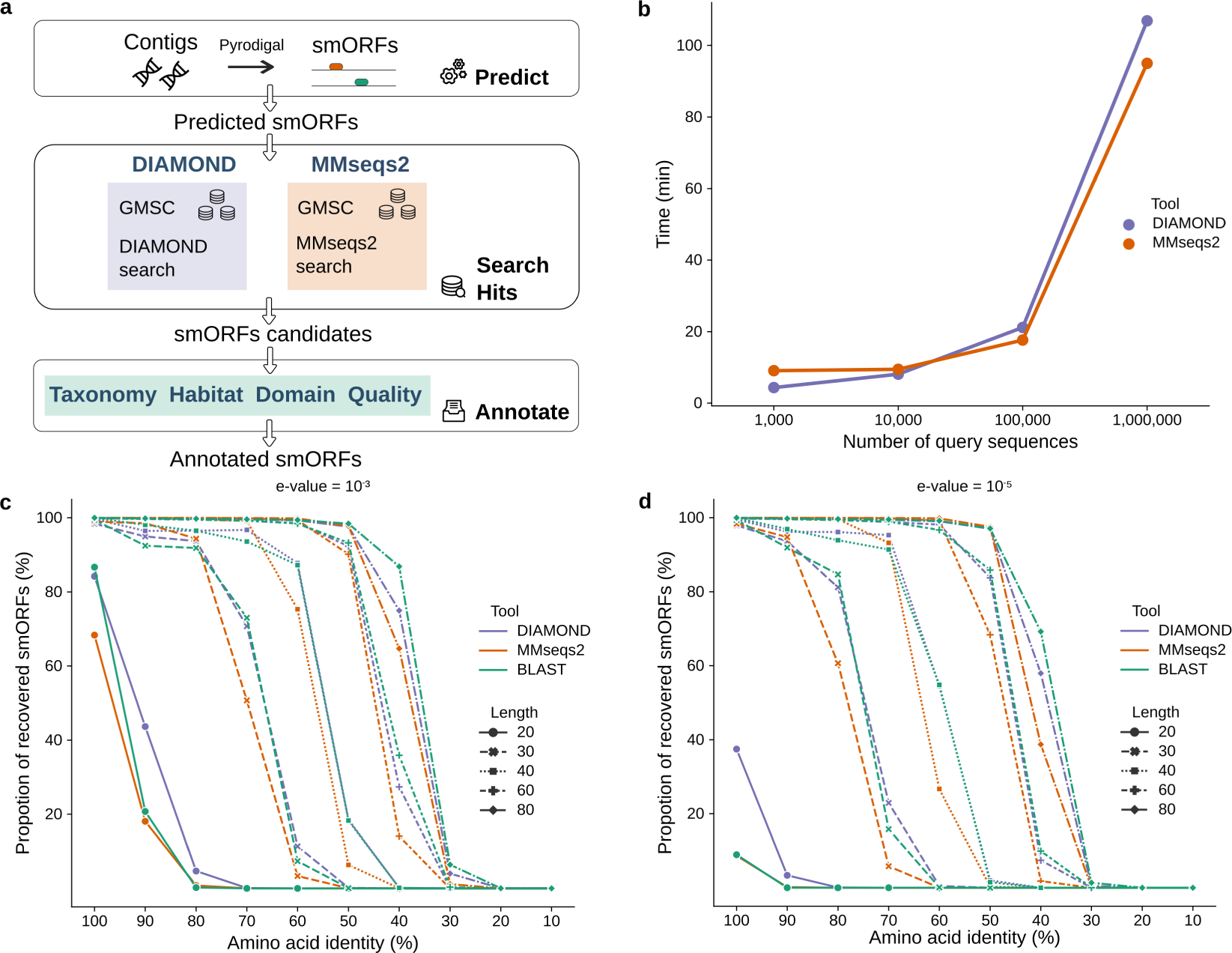
Workflow and benchmark of GMSC-mapper. **(a)** GMSC-mapper uses Pyrodigal to predict small proteins with < 100 amino acids from contigs. Users can alternatively provide smORF or protein sequences directly, skipping the initial step of gene prediction. DIAMOND or MMseqs2 are used for finding homologues within GMSC. In the end, GMSC-mapper combines all alignment hits and provides detailed annotations of small proteins. **(b)** Time cost tests were performed among different numbers of input sequences from 1,000 to 1,000,000 using DIAMOND and MMseqs2 (Methods). We compared the number of recovered sequences with different lengths (20, 30, 40, 60, and 80 amino acids) at different amino acid identities from 10% to 100% using DIAMOND, MMseqs2, and BLAST (Methods). The recovered number is influenced by the E-value cutoff used (10*^−^* ^3^ in **(c)** and 10*^−^* ^5^ in **(d)**).

GMSC-mapper performs *de novo* prediction and annotation of small proteins encoded by smORFs in user-provided genomes or assembled metagenomes (Methods). For this, it first uses Pyrodigal^35,52^ to predict small proteins from assembled contigs and then it uses DIAMOND^53^ or MMseqs2^54^ to align these predictions against the GMSC. To minimize computational resource usage, GMSC-mapper only searches family representatives, but it returns the set of matching smORFs and the annotation of the matches (*e.g.*, habitat and taxonomy) as well as links to GMSC identifiers.

We compared DIAMOND to MMseqs2 for this task and observed that DIAMOND is faster than MMseqs2 when the number of query sequences is below 10,000, while MMseqs2 is slightly faster than DIAMOND when the number of queries is above 10,000 (Fig. 5b). In addition, we compared the number of recovered sequences (Fig. 5c) with either of these tools or BLAST^55^, by randomly modifying sequences in the catalogue and aligning these modified versions back to the catalogue of family representatives. All three tools can find a high-identity match if it is present in the database. With increasing sequence size, these tools can match more distant homologous sequences. In this case, DIAMOND achieves almost the same sensitivity as BLAST and is superior to MMseqs2^56^.

However, independently of the method used, when the sequences are too short (20 amino acids), the rate of recovery decreases drastically. Fundamentally, for short sequences in a large database, even an identical match has a high likelihood of arising by chance^57,58^. This will manifest itself in a high e-value^59^ for true positives, making it impossible to distinguish false and true positive matches based on sequence comparisons alone. Therefore, while the use of a higher e-value threshold will recover a larger fraction of true matches (> 80% recovered with DIAMOND using 10*^−^* ^3^ compared to circa 40% using 10*^−^* ^5^, see Fig. 5c-d), the false discovery rate (FDR) will also increase^57^.

## Discussion

Here, we constructed the global microbial smORFs catalogue (GMSCv1, in its first version), which contains approximately 1 billion smORF sequences, of which 43 million are *high-quality predictions*, representing a large increase in the number of smORF sequences previously reported and serving as a resource for the microbiome research community. Previously, most of the widely studied microbial small proteins were accidentally discovered in isolated and cultured bacterial species^5^. The large-scale discovery of small proteins has made great progress in recent years. Sberro et al. conducted the characterization of conserved small proteins in the human microbiome, revealing their potential various functions^33^. In our work, we have expanded the discovery of small proteins to 75 distinct habitats worldwide. In our catalogue, only a small fraction are homologous to reference small protein datasets with the vast majority of the novel small proteins being found in non-human-associated habitats (Supplementary Fig. 4b). On the other hand, it encompasses most of the sequences in other resources. For each smORF and small protein family, we provide comprehensive annotations, including taxonomy, habitats, and conserved domains.

A major difficulty in finding biologically-functional smORFs is that false positive predictions are common. One of the underlying principles in our efforts is that finding identical or highly similar sequences in multiple samples increases the likelihood of a true prediction. Therefore, we discarded singleton predictions in our data. This principle also underlies the GMSC-mapper tool, which enables users to find matches from their datasets in GMSCv1.

As previously done^33^, we have only conducted RNAcode^37^ on small protein families with at least 8 members to identify smORFs families with transcription signatures. This approach may, however, fail to identify some rapidly evolving functional smORFs. In addition, given the limited size and number of existing datasets of metatranscriptomes, (meta)Ribo-Seq and metaproteomes, the *high-quality predictions* are expected to underestimate the true diversity.

Computing approaches and concepts developed over decades for longer proteins do not necessarily work well for small sequences. For example, for the alignment of very short sequences, the minimum achievable E-value will be lower bounded^59^. Even an identical match will obtain a relatively high e-value as short identical matches can occur by chance. Furthermore, traditional databases lack small proteins, so that functional assignment by orthology or with HMMs only returns a prediction for a minute fraction of all small proteins. We lack functional predictions for most small proteins in our dataset, even for those small protein families that are ubiquitous. Similarly, tools for predicting whether proteins are transmembrane or secreted are not optimized for small proteins and our results should be interpreted in this context. Considering the above, in related work, we used machine learning^36^ to identify candidate antimicrobial peptides (AMPs) from the GMSC^45^. However, functional prediction for small proteins remains an open challenge, open to new approaches. Overall, our resource shows the immense and underexplored diversity of small proteins across different habitats and taxonomy, and highlights the gaps in our scientific knowledge, while constituting a resource for the research community.

## Methods

### Collection of global metagenomes and high-quality microbial genomes and prediction of smORFs

In total, 63,410 publicly-available global assembled metagenomes from the SPIRE database^21^ collection were used, having been processed as previously described. Briefly, publicly available data (as of 1 January, 2020) were downloaded from the European Nucleotide Archive and short reads that were at least 60 bps after trimming positions with quality < 25^60^ were assembled into contigs using MEGAHIT 1.2.9^61^. Additionally, we downloaded 87,920 high-quality isolate microbial genomes from the ProGenomes2 database^34^.

We then used the modified version of Prodigal^35^ in Macrel 0.5^36^ to predict open reading frames (ORFs) ≥ 30 base pairs (bps) on the assembled contigs as well as those from Progenomes2 database. The ORFs encoding small proteins (here defined as those up to 100 amino acids) were considered smORFs. In total, 4,599,187,424 smORFs were predicted, of which 99.25% (4,564,570,019) originated in metagenomes and 0.75% (34,617,405) originated in microbial genomes of the Progenomes2 database.

We recorded the habitats of smORFs according to their source samples using the habitat microontology introduced in SPIRE database^21^. We further grouped the habitats into 8 broad categories: mammal gut, anthropogenic, other-human, other-animal, aquatic, human gut, soil/plant, and other.

### Non-redundant smORFs catalogue construction and method validation

All the smORFs were first deduplicated at 100% amino acid identity and 100% coverage. The number of smORFs was reduced from 4,599,187,424 to 2,724,621,233, with 2,307,007,717 singletons that occurred only once and 417,613,516 non-singletons.

We hierarchically clustered the non-singletons at 90% amino acid identity and 90% coverage, resulting in 287,926,875 clusters, which we will henceforth refer to as families, using Linclust^54,62^ with the following parameters: -c 0.9, –min-seq-id 0.9.

Of these clusters, 47.5% contain a single sequence (singleton clusters). To rule out the possibility that this was due to the fact that Linclust^54,62^ is a heuristic method that is not specifically designed for short sequences, we aligned a randomly selected 1,000 singleton clusters against the representative sequences of non-singleton clusters (*i.e.*, those containing ≥ 2 sequences) using SWIPE^63^ with the following parameters: -a 18 -m ‘8 std qcovs’ -p 1. The alignment threshold was E-value < 10*^−^* ^5^, identity ≥ 90%, and coverage ≥ 90%. Only 44 (4.4%) of the randomly selected sequences had homologues not identified by Linclust^54,62^ (Supplementary Fig. 1a).

In addition, to test whether the sequences of clusters are significant at the 90% identity level. As described above, 1,000 sequences were randomly selected and aligned against the representative sequences of their clusters using SWIPE^63^. Each pair of sequences was considered significant when E-value < 10*^−^* ^5^, identity ≥ 90%, and coverage ≥ 90%. Results showed that 99.2% (992 / 1,000) of sequences in clusters are clustered significantly (Supplementary Fig. 1b).

When clustering, we initially discarded the 2,307,007,717 singletons because singletons are enriched in artifactual smORFs^36^. However, we considered that singletons that are homologous to larger clusters should not be discarded as the homology itself provides further evidence of biological relevance. Therefore, we aligned singletons to the representative sequences of clusters with 90% sequence identity and 90% coverage using DIAMOND^53^ using parameters: -e 10*^−^* ^5^ –id 90 -b 12 -c 1 –query-cover 90 –subject-cover 90. This identified homologues for 547,356,980 sequences. Combining these with the 417,613,516 non-singleton sequences identified earlier resulted in a catalogue of 964,970,496 sequences which we termed the *smORFs catalogue*.

### Sample-based smORFs rarefaction curves

Samples were randomly permuted 24 times to calculate the total number of non-redundant smORFs captured as the number of samples increased. We took the average across the permutations as the final estimate.

### Quality control of the GMSC

We conducted several *in silico* quality tests and matched genomic predictions to other publicly-available experimental data.

A smORF predicted at the start of a contig that is not preceded by an in-frame STOP codon risks being a false positive originating from an interrupted fragment. Therefore, we checked for the presence of an upstream in-frame STOP and found one in 41.93% of GMSC smORFs. For other smORFs, however, we could not determine whether there were other genes present upstream of them.

To avoid spurious smORFs, we used HMMSearch^64^ with the `--cut_ga` option to search smORFs against the AntiFam database^65^, which contains a series of confirmed spurious protein families. It indicated that 99.8% of GMSC smORFs are not from any known false-positive protein families.

We used RNAcode^37^, a tool to predict the coding potential of sequences based on evolutionary signatures, to identify the coding potential of 25,744,932 smORF families containing ≥ 8 sequences. It showed that 6.3% of the smORF families, including 32.1% of GMSC smORFs have a coding potential with p-value < 0.05.

Furthermore, we searched for evidence that these smORFs are transcribed and/or translated. For this step, we downloaded 221 publicly available metatranscriptomic datasets from the NCBI database paired with the metagenomic samples we used in our catalogue (Supplementary Table 3). These samples are from the human gut, peat, plant, and symbionts. To keep the procedure computationally feasible, we mapped reads against the representative sequences of smORF families by BWA^66^. Then we used NGLess^60^ with ‘unique_only’ for the ‘multiple’ argument of the count built-in function to only count uniquely mapped inserts. A smORF family was considered to have transcriptional evidence if its representative has reads mapped to it in at least 2 samples. The results showed that 34.3% of the smORF families including 38.2% of the GMSC smORFs have transcriptional evidence.

We downloaded 142 publicly available Ribo-Seq datasets from the NCBI database (Supplementary Table 3). We also mapped reads against representative sequences of smORF families by BWA^66^. Then we used NGLess^60^ with ‘unique_only’ for the ‘multiple’ argument of the count built-in function to only count uniquely mapped inserts. A smORF family was considered to have translation evidence only if its representative has reads mapped to it in at least 2 samples. The results showed that 0.7% of the smORF families including 1.2% of the GMSC smORFs have translational evidence.

Moreover, we downloaded peptide datasets from 108 metaproteomic projects from the PRIDE database^67^ (Supplementary Table 3). We matched GMSC smORFs to the identified peptides of each project. If the total k-mer coverage of peptides on a smORF is greater than 50%, then the smORF is considered translated and detected, as in a previous study^68^. In total, 1.3% of the GMSC smORFs were matched.

Sequences that passed all *in silico* tests above as well as matching transcriptional or translational data were regarded as *high-quality predictions* and, in total, 43,617,914 (4.5%) of the 964,970,496 GMSC smORFs fulfilled these criteria.

### Comparison with reference small protein datasets

We downloaded bacterial and archaeal protein sequences from RefSeq in March 2023^38^. The sequences with fewer than 100 amino acids are considered small proteins, and redundancy was subsequently removed with 100% amino acid identity and 100% coverage. A total of 16,333,323 non-redundant bacterial small proteins and 368,769 non-redundant archaeal small proteins from RefSeq were included in the comparison. We also included the 444,053 small protein clusters and 4,539 conserved small protein families from Sberro’s human microbiome study^33^. We compared our *smORFs catalogue* to these reference datasets using DIAMOND with the ‘–more-sensitive’ mode, retaining significant hits (E-value < 10*^−^* ^5^).

### Conserved domain annotation

We downloaded the Conserved domain database (CDD)^39^ from ftp://ftp.ncbi.nih.gov/pub/mmdb/cdd/little_endian/Cdd_LE.tar.gz in September 2022, which contains models from Conserved Domains (CD) curated at NCBI, Pfam^44^, SMART^69^, COGs^47^, PRK^70^, and TIGRFAMs^71^. All the representative sequences of small protein families and randomly selected 10,000 prokaryotic proteins from the global microbial gene catalogue v1.0^20^ were searched against the Conserved domain database by RPS-BLAST^40,41^. Hits with an E-value maximum of 0.01 and at least 80% of coverage of PSSM’s length were retained and considered significant. Pfam accessions were grouped by Pfam clan^72^ or the first phrase before the comma in their short description.

### Taxonomic annotation and taxonomic breadth analysis

The taxonomy of assembled contigs encoding the small proteins was annotated using MMseqs2 taxonomy^42^ against the GTDB database^73^ release r95. However, in figures and text, we used updated taxa names (e.g., Bacillota instead of Firmicutes). We characterized the taxonomy of predicted smORFs based on the taxonomy of contigs and microbial genomes^34^ from which the smORFs were predicted. We subsequently assigned taxonomy for GMSC smORFs and families using the lowest common ancestor (LCA), ignoring the un-assigned ranks to make them more specific.

The small protein families with at least three members were subsequently used to perform taxonomic breadth analysis. Each family was classified according to *(i)* whether it is single or multi-habitat; *(ii)* whether it is single or multi-genus; and *(iii)* whether it is annotated with a Pfam domain^44^. Multi-genus families are more common in multiple habitats than the entire families (61.8% vs. 52.0%; P-value < 10*^−^* ^308^, hypergeometric test). Multi-genus families are annotated with Pfam domains at a higher rate (9.91% vs 7.52%; P-value < 10*^−^* ^308^, hypergeometric test). As these results could have been confounded by differences in family size distributions, we randomly downsampled the data to keep the same number of families at each size between multi-genus families and the whole families. In that case (as presented in the main text), the difference was 61.8% vs. 57.5% (P-value < 10*^−^* ^308^, hypergeometric test) for the proportion of families in multiple habitats and 9.91% vs. 8.15% (P-value < 10*^−^* ^308^, hypergeometric test) for the proportion of Pfam annotated families.

### Density calculation

The density of smORFs was defined as ρ = *n_sm O RF s_* / L, where *n_sm O RF s_* is the number of redundant smORFs and L is the assembled megabase pairs (Mbps)^32,45^. The density was calculated by summing all assembled base pairs for contigs assigned to each taxonomic rank. We assume a scenario where the starting positions of smORFs in an assembled large contig are independent and unifo<u>rmly rando</u>m. Therefore, the standard sample proportion error was calculated as 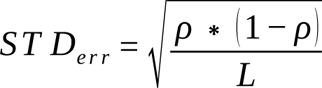 and was used to calculate the margin of error at a 95% confidence interval (Z = 1.96). We did not further consider the calculated values with a margin of error >10%.

### Cellular localization prediction

To detect potential transmembrane proteins, we ran TMHMM-2.0^48^ on the representative sequences of small protein families. Then, to identify potentially secreted small proteins, we used SignalP-5.0^49^ on the representative sequences of small protein families. For families classified as archaea, we used ‘-org arch’, while for the others we combined the outputs of ‘-org gram+’ and ‘-org gram-’ modes.

### Construction and evaluation of GMSC-mapper

GMSC-mapper supports assembled contigs, smORF sequences, or protein sequences as inputs. It uses Pyrodigal^35,52^, which is a faster implementation of the Prodigal algorithm, to predict ORFs potentially coding for small proteins (those with fewer than 100 amino acids) from contigs. Gene prediction is skipped when inputs are smORF or protein sequences. Then DIAMOND^53^ or MMseqs2^54^ are used for homologous alignment against GMSC. Finally, it combines all the alignment hits information and provides detailed annotation of small proteins.

To determine the optimal default sensitivity mode, we tested different sensitivity parameters for DIAMOND and MMseqs2. We aligned 10,000 randomly selected sequences back to the family representatives and counted the number of recovered sequences while monitoring the computational time. We use the “–sensitive” mode as the default sensitivity parameter for DIAMOND, which provides the best balance between sensitivity and speed. The use of more sensitive modes resulted in little or almost no increase in the number of recovered sequences, but a substantial increase in time usage. For MMseqs2, we keep the original default sensitivity parameter (5.7) considering that the number of recovered sequences and the time both increase with the increase of sensitivity (Supplementary Fig. 5a-d).

We then tested the time costs among different numbers of input sequences using the “– sensitive” mode of DIAMOND and the default sensitivity parameter (5.7) of MMseqs2. GMSC-mapper can annotate 100,000 input sequences in approximately 20 minutes with 20 threads.

Furthermore, we compared the number of recovered sequences with different identities using different alignment tools. We randomly selected and mutated 10,000 sequences of different lengths (20, 30, 40, 60, and 80) from the family representatives, with different identities from 10% to 90%. We aligned them back to the family representatives using DIAMOND, MMseqs2, and BLAST^55^, respectively. When the query sequence and the target sequence are the same, we consider them as the recovered sequences.

Timing measurements were performed using a server equipped with an AMD EPYC 7763 64-Core processor and 2TB of RAM memory.

### Statistical analyses

Statistical analyses was carried out in Python, using Pandas^74^, NumPy^75^, and SciPy^76^.

### GMSC web resource

GMSC webserver is hosted at the address https://gmsc.big-data-biology.org, where an implementation of GMSC-mapper can be accessed. The website implementation is based on Elm-Lang. The API implementation is based on Python.

## Data Availability

Global metagenomic data are publicly available at the European Nucleotide Archives (ENA). The accession numbers for samples and studies are listed in Supplementary Table 1. Microbial genomes are publicly available in the Progenomes2 database.

The global microbial smORFs catalogue (GMSC) resource has been deposited in Zenodo under DOI: 10.5281/zenodo.7944370. The resource is freely available at https://gmsc.big-data-biology.org. Users can query small protein sequences by using GMSC-mapper through the web interface or select their interesting small proteins by habitats and taxonomy.

## Code Availability

The codes used to generate and analyse the global microbial smORFs catalogue (GMSC) are available at https://github.com/BigDataBiology/Duan2023GMSCv1_Construction_And_Analysis. GMSC-mapper is open source and at https://github.com/BigDataBiology/GMSC-mapper.

## Acknowledgements

This work was supported by National Key R&D Program of China (2020YFA0712403, 2018YFC0910500), National Natural Science Foundation of China (61932008, 61772368, T2225015), Shanghai Science and Technology Innovation Fund (19511101404), Shanghai Municipal Science and Technology Major Project (2018SHZDZX01), Greater Bay Area Institute of Precision Medicine (Guangzhou) (Grant No. IPM21C008), and the Australian Research Council (grant FT230100724).

We thank Ben Woodcroft (Queensland University of Technology) for helpful comments on a previous version of the manuscript.

## Author contributions

LPC conceptualized and designed the study. YD, CDSJ, TSBS, AF, LPC, MK curated the data. YD, CDSJ, BLSD, and CZ analysed and visualized the data. LPC, XMZ, and PB supervised the project. YD and LPC wrote the original draft. All authors reviewed and contributed to the manuscript.

## Supplementary Figures

**Supplementary Fig. 1.**
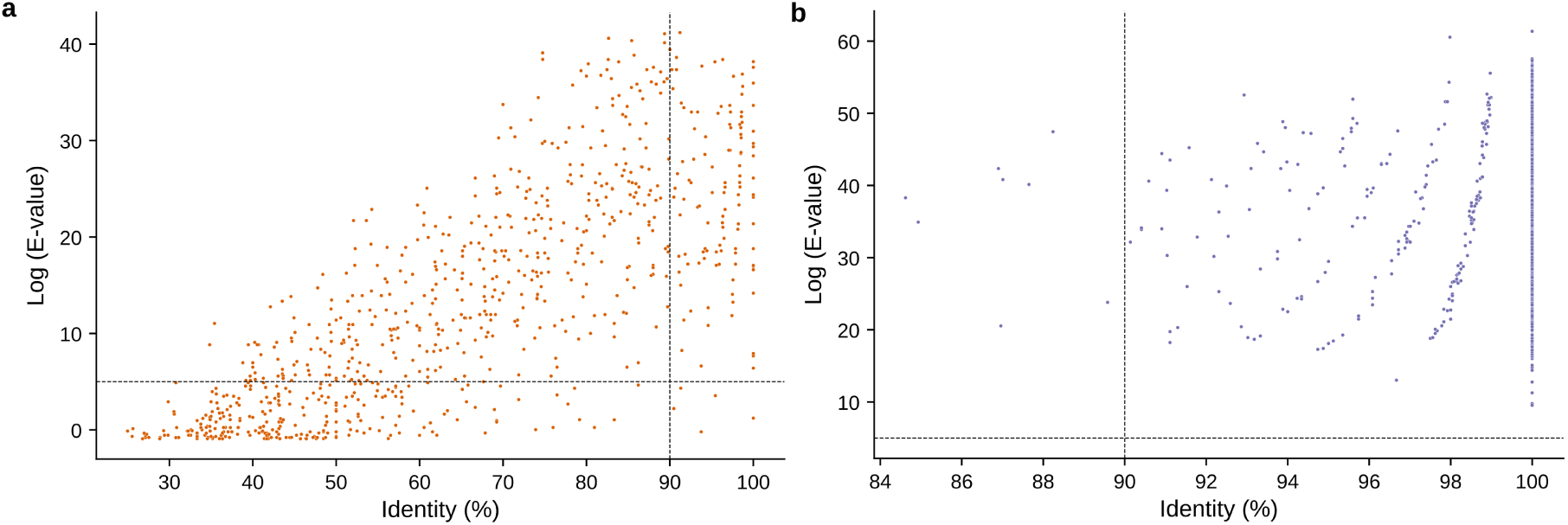
Clusters significance validation. **(a)** We aligned a randomly selected 1,000 singleton clusters against the representative sequences of non-singleton clusters using SWIPE (an exhaustive search method). The hits with E-value ≤ 10*^−^* ^5^, identity ≥ 90%, and coverage ≥ 90% were considered significant and represent false negatives from the heuristic alignment method used. **(b)** We aligned a randomly selected 1,000 sequences against the representative sequences of clusters they originate from using SWIPE. The hits with E-value ≤ 10*^−^* ^5^, identity ≥ 90%, and coverage ≥ 90% were considered significant.

**Supplementary Fig. 2.**
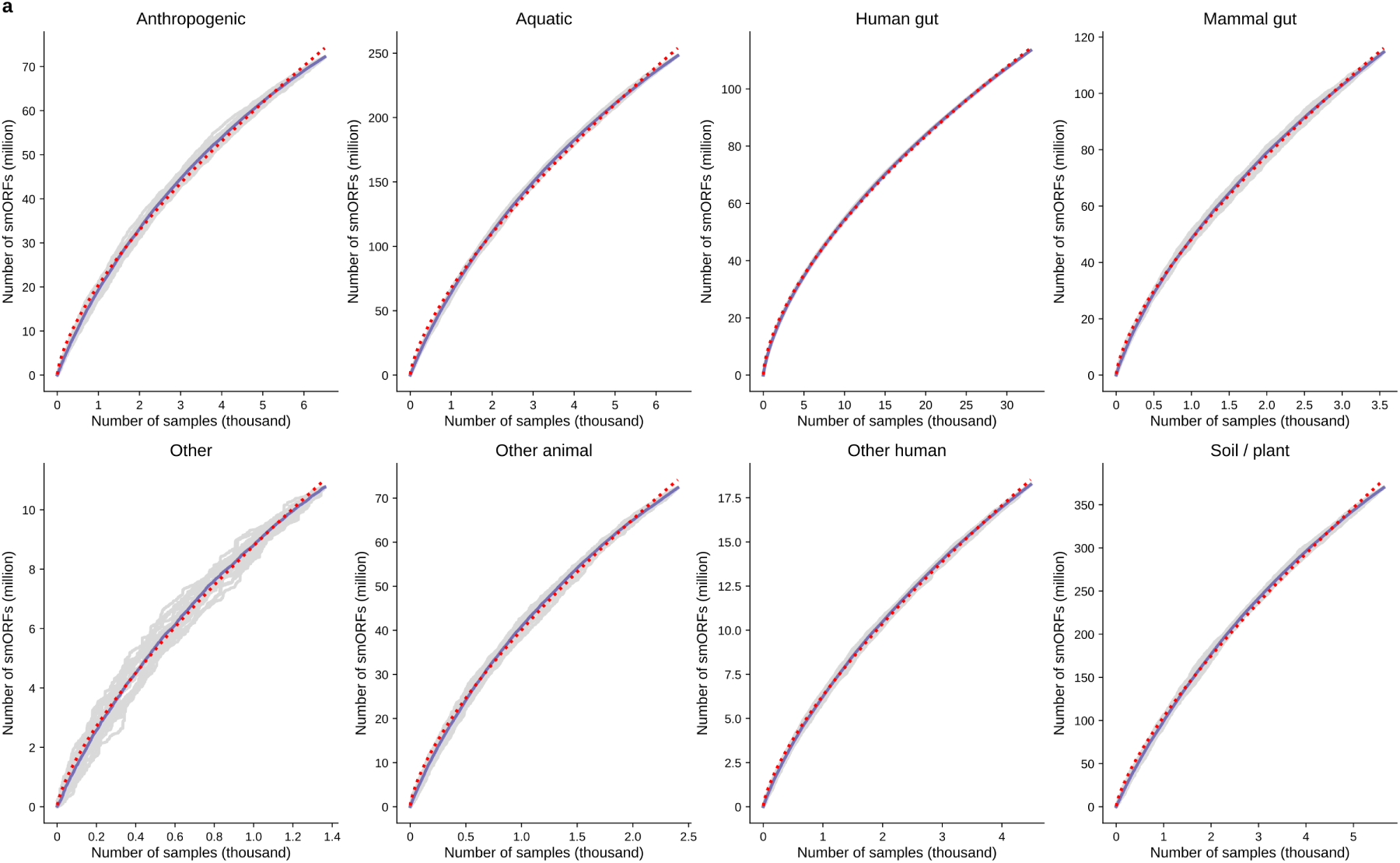
The smORF accumulation curves across habitats. **(a)** The grey lines represent the 24 permutations of sample selection for each broad habitat category. The blue line is the average of these permutations. The red line is the fit of Heap’s Law (*N* =*k · s a m p l e^al^ ^ph^ ^a^*). It indicates that the amount of smORFs in any habitat is not saturated.

**Supplementary Fig. 3.**
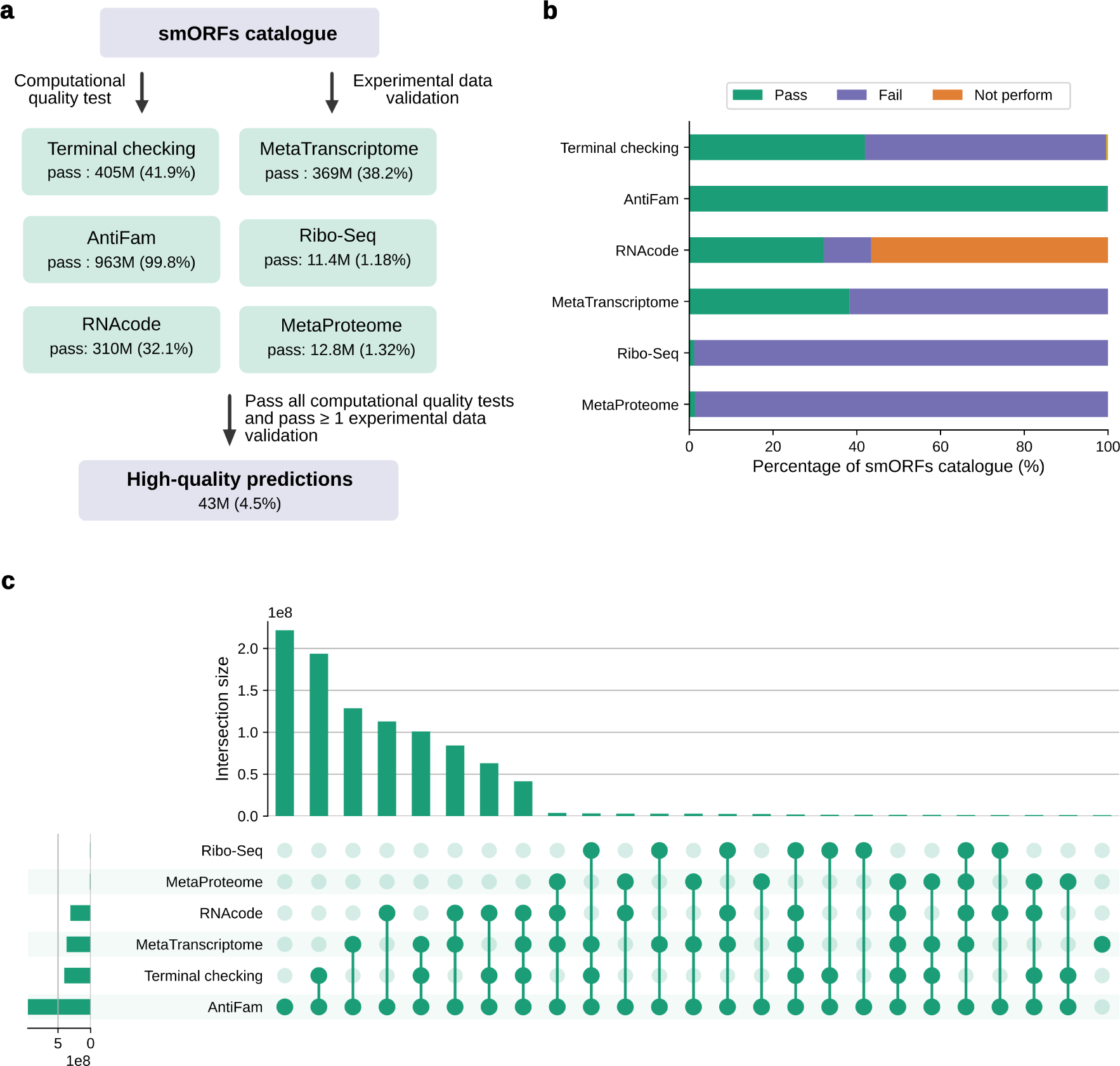
Quality assessment workflow and overlap. **(a)** The computational quality tests include: (i) terminal checking to estimate the risk of false positives that the smORF is derived from interrupted fragments; (ii) AntiFam searches to avoid spurious protein families; and (iii) RNAcode estimated coding potential. The experimental data validation consists of mapping the metatranscriptomic and Ribo-Seq reads downloaded from the public database and exactly matching metaproteomic peptides downloaded from the Proteomics Identification Database (PRIDE). SmORFs were considered high-quality predictions if they passed all computational quality tests and were found in at least one experimental dataset. **(b)** Fraction of GMSC smORFs each test. RNAcode was performed only on clusters with at least 8 members. Terminal checking was performed only on smORFs derived from metagenomes. **(c)** The upset plot shows the number of overlapping sequences passing each quality testing method.

**Supplementary Fig. 4.**
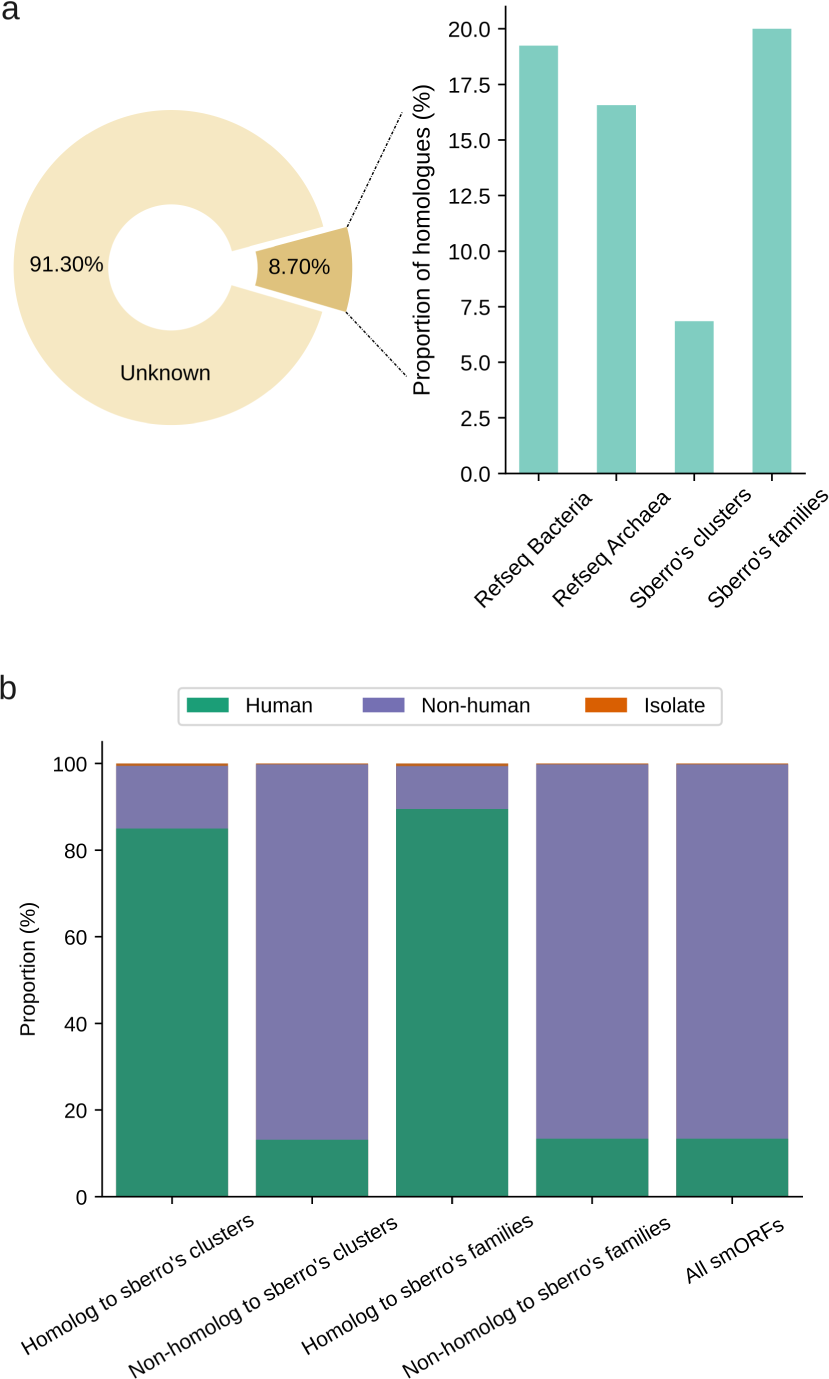
Comparison of reference small protein datasets. **(a)** Shown is the fraction of smORFs from high-quality predictions that are homologous to reference small protein datasets. **(b)** The comparison of the proportions of smORFs from human or non-human habitats between homologues or non-homologues to small protein clusters and conserved families from Sberro human microbiome dataset.

**Supplementary Fig. 5.**
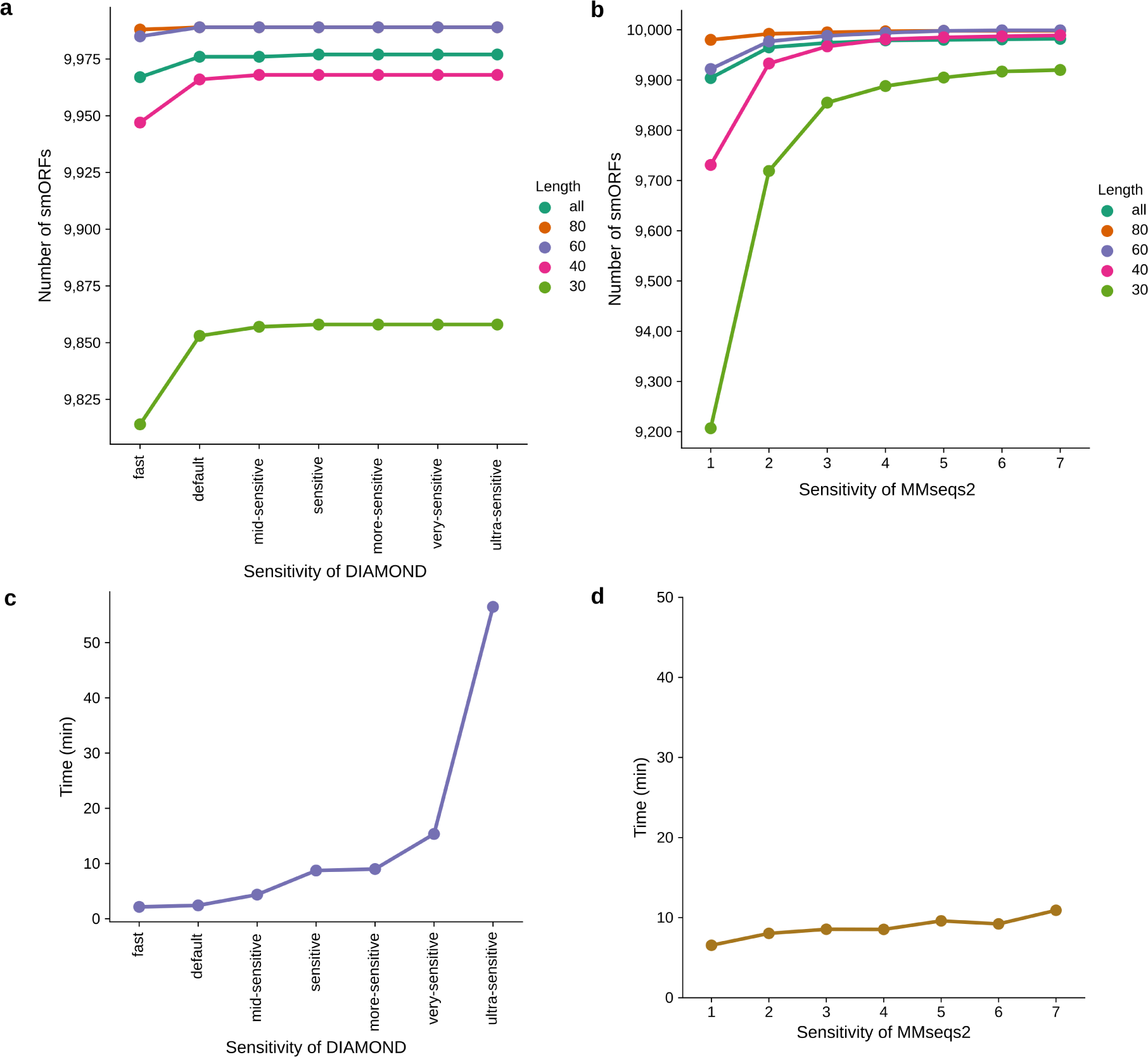
Benchmark of sensitivity modes between DIAMOND and MMseqs2. **(a)** Shown is the recovery amount of 10,000 randomly selected smORFs with different lengths under different sensitivity modes by DIAMOND. **(b)** Shown is the recovery amount of 10,000 randomly selected smORFs with different lengths under different sensitivity parameters by MMseqs2. **(c)** Shown is the time cost for DIAMOND to map 10,000 randomly selected smORFs to the smORF family representatives under different sensitivity modes. **(d)** Shown is the time cost for MMseqs2 to map 10,000 randomly selected smORFs to the smORF family representatives under different sensitivity parameters.

## Supplementary Table Legends

**Supplementary Table 1 Metadata of metagenomes and isolated genomes** Each metagenomic sample is provided with project accession, publication, habitat including environment and host, geographic coordinates, and collection dates. Basic assembly information for each sample includes the number of assembled base pairs (bps) and the assembly N50. The number of non-redundant *smORFs catalogue* and the number of smORF families for each metagenomic sample and the isolated genomes from the Progenomes2 database are shown.

**Supplementary Table 2 The distribution of smORFs in the broad habitat categories** Displayed the various habitats included in each broad habitat category, number of samples, number of redundant smORFs, number of non-redundant smORFs catalogue, and number of smORF families.

**Supplementary Table 3 Metatranscriptomic, Ribo-Seq, and metaproteomic datasets used in the experimental validation** We downloaded 221 publicly available metatranscriptomic datasets, which are paired with the metagenomic samples that we used in the catalogue. We downloaded 142 publicly available Ribo-Seq datasets. We downloaded peptide datasets of 108 metaproteome projects from the Proteomics Identification Database (PRIDE). The accession and source of the projects are listed.

**Supplementary Table 4 Conserved domain annotation of the catalogue** The number of annotated sequences and detailed descriptions for each conserved domain are provided. Pfam accessions are grouped by Pfam clan and short description.

**Supplementary Table 5 Conserved domain annotation of the small protein families present in multiple phyla and habitats** The number of members of the small protein families, the number of species of the small protein families, and conserved domain annotation with description are provided.

**Supplementary Table 6 smORF density across different phyla** The number of redundant smORFs and the assembled base pairs are shown and used to calculate the smORF density for each phylum.

